# Muscle and intestine innexins with muscle Deg/Enac channels promote muscle coordination and embryo elongation

**DOI:** 10.1101/2024.07.12.603207

**Authors:** Flora Llense, Teresa Ferraro, Xinyi Yang, Hanla Song, Michel Labouesse

## Abstract

Body axis elongation represents a fundamental morphogenetic process in development, which involves cell shape changes powered by mechanical forces. How mechanically interconnected tissues coordinate in organismal development remains largely unexplored. During *C. elegans* elongation, cyclic forces generated by muscle contractions induce remodeling of adherens junctions and the actin cytoskeleton in the epidermis, facilitating gradual embryo lengthening. While previous studies have identified key players in epidermal cells, understanding how muscle cells coordinate their activity for proper embryo elongation remains unsolved. Using a Calcium sensor to monitor muscle activity during elongation, we identified two cells in each muscle quadrant with a leader cell function that orchestrate muscle activity within their respective quadrants. Strikingly, ablation of these cells halted muscle contractions and delayed elongation. A targeted RNAi screen focusing on communication channels identified two innexins and two Deg channels regulating muscle activity, which proved required for normal embryonic elongation. Interestingly, one innexin exhibits specific expression in intestinal cells. Our findings provide novel insights into how embryonic body wall muscles coordinate their activity and how interconnected tissues ensure proper morphogenesis.

## Introduction

Tissue and embryo morphogenesis rely on intermingled cellular processes regulated by biochemical signalling pathways, and physical forces from surrounding cells and tissues (Zinner, and Liberali, 2020). Most organs consist of multiple tissue layers that are mechanically coupled via the extracellular matrix (Nelson and Gleghorn, 2012; Mitchell *et al*., 2022). Understanding how cell shape changes influence embryo-scale dynamics remains challenging. While some molecular players in mechanical force transmission are known, coordination between mechanically coupled tissues remains unclear.

*C. elegans* embryo elongation, which depends on cell shape changes driven by the epidermis, provides an ideal system to tackle these questions (Vuong-Brender, *et al*., 2016). It involves two phases: the first, from the comma to the 2- fold stages, depends on tension and stiffness anisotropy in the epidermis (Vuong-Brender *et al*., 2017); beyond the 2-fold stage, it requires muscles, which are tightly attached to the epidermis through hemidesmosomes and a common extracellular matrix (Fig. 1A) (Vuong-Brender, *et al*, 2016). Four body wall muscle quadrants are running along the anterior-posterior direction, two dorsal and two ventral (Gieseler, *et al*, 2017). Muscle contractions trigger a mechanotransduction pathway involving the p21-activated kinase PAK-1 within the overlying epithelial layer, which triggers hemidesmosome remodelling and actin cytoskeleton reorganization (Zhang *et al*., 2011; Lardennois *et al*., 2019).

**Figure 1.**
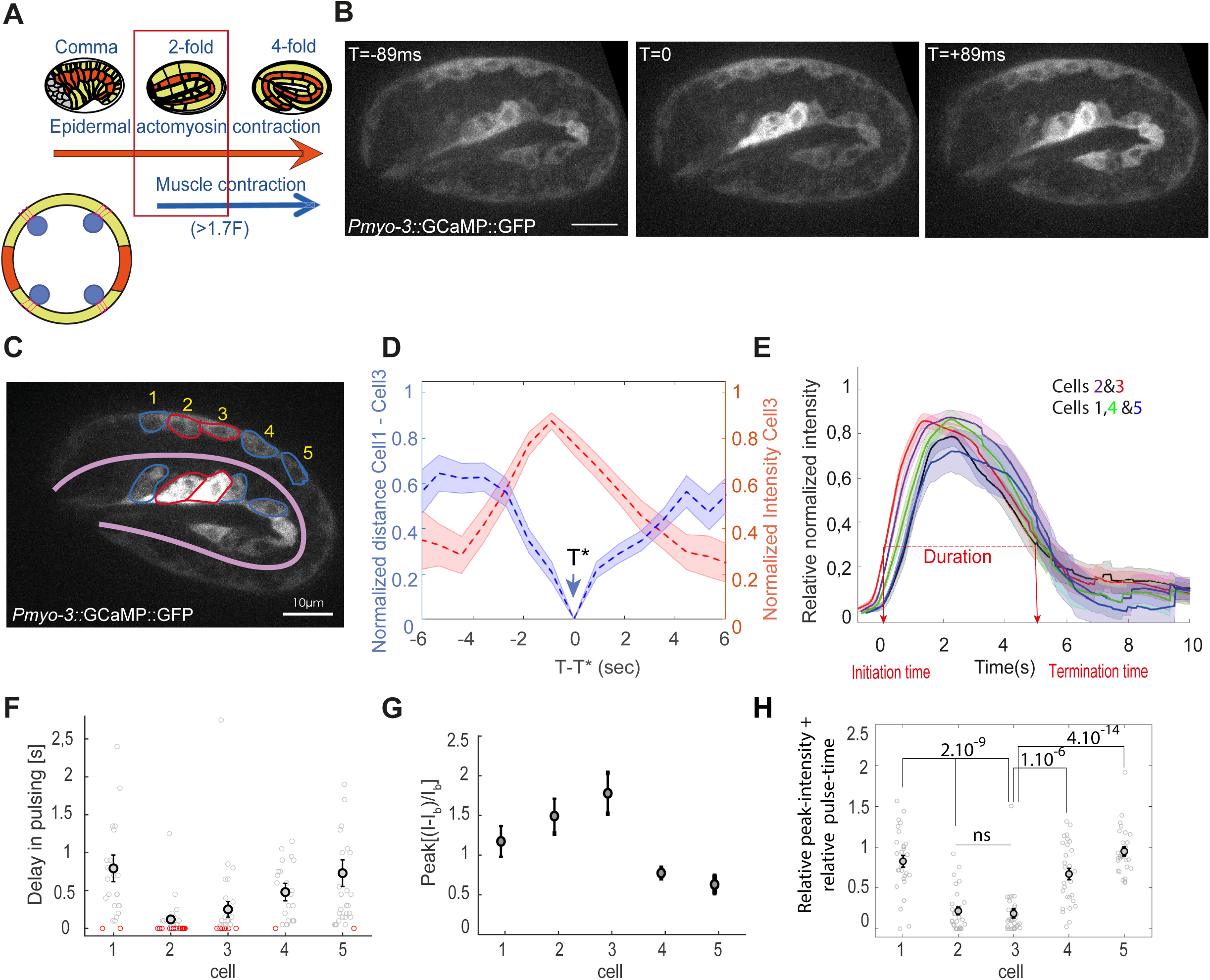
Embryonic muscle activity. **(A)** Scheme of elongation depicting the lateral (orange) and dorso-ventral epidermal cells (green). Red rectangle, time focus of the study. The cross-section below shows the relative positions of muscle cells (blue) just underneath the dorso-ventral epidermis, with the hemidesmosomes (red lines). **(B)** Snapshots from a timelapse showing muscle activity in an embryo labelled with GCaMP. T=0, time of the highest intensity. **(C)** Image of a two-fold embryo showing the GCaMP in the five cells with the highest fluorescence levels within in each muscle quadrant. **(D)** Time correlation between muscle contractions (cell-1-3 distance, blue), and the normalized GCaMP intensity of cell-3 (red). T*, time of minimal distance between cell-1/3 (blue arrowhead). the calcium peak occurs 0.89 s before the contraction peak. **(E)** Graph showing the calcium wave starting in cells-2/3 and propagating to cells-1/4/5. In (D-E), lines, average; shadow areas, standard error of the mean (SEM). **(F)** Dot-plot showing the delay between Ca^++^ pulses among the five cells-1/5; the cell pulsing first is in red for a given wave. **(G)**. Normalized GCaMP maximum intensity relative to the b baseline fluorescence (I_b_) for each cell. I_b_ is chosen as the lowest intensity value among the 5 cells in one wave. **(H)** Plot showing the relative peak intensity and relative pulse time combined together (see methods).

Whereas muscle cell dynamics in the adult and larval stages are well known (Gieseler, *et al*, 2017), little is known about embryonic muscle dynamics. In particular, we do not know how muscle cells initiate their contractions in embryos and how they coordinate their activity within each quadrant. The regulation of muscle contractions is unlikely to depend on synaptic input. Indeed, axonogenesis is still underway during the first embryonic contractions (Wu *et al*., 2011; Shah *et al*., 2017; Moyle *et al*., 2021). Moreover, mutants defective for acetylcholine or GABA release such as *unc-17* or *snb-1*, which cause complete paralysis in larvae, do not trigger the Paralyzed at two-fold (Pat) typical of mutants affecting myofilament attachment (such as *pat-2/3*) (Alfonso *et al*., 1993; Nonet *et al*., 1998)

Here, to improve our understanding of muscle coordination in elongating embryos, we explored muscle activity by calcium imaging. To identify new molecular players involved in regulating muscle activity, we performed a RNAi screen targeting proteins involved in cell-cell communication. Our work thus further explains how muscles and the epidermis jointly drive embryonic morphogenesis.

## Results and discussion

### Embryonic body muscles organization and dynamics

We aimed to investigate how muscle activity is initiated and controlled during elongation (Fig. 1A). To track muscle activity over time, we used the ratiometric GCaMP calcium sensor expressed under the muscle-specific *myo-3* promotor (Schwarz, *et al*, 2012). A time-lapse analysis was conducted to examine the variation of probe intensity in two lateral muscle quadrants within the field of view (Fig. 1B, Fig. S1A). We first observed mini-spikes occurring in a random pattern in all muscles before the first muscle contraction in dorsal quadrants (Movie1), as also reported elsewhere (Ardiel *et al*., 2017). Subsequently, the first contractions coincided with a brighter GCaMP signal in five adjacent cells in each quadrant (Movie1), whereas it became dimmer in other cells. We also noticed that dorsal and ventral quadrants were not synchronized. Interestingly, our recent lightsheet microscopy analysis of adherens junction extension revealed that epidermal deformations start in this area (Yang *et al*., 2024). We called these cells cell-1 to cell-5 according to their positions along the anterior-posterior axis (Fig. 1C). Cell-2 and cell-3 were located posterior to pharyngeal PHA-4 positive cells, and cell-4 roughly at the anterior-posterior level of the V1 lateral epidermal cell (Fig. S1B and C). We conclude that the identity of the left cell-2 and cell-3, which play a key role (see below), are dorsally MSapppap (cell-2d) and Daaap (cell-3d); and ventrally Daaaa (cell-2v) and Dapaa (cell- 3v).

We analyzed the calcium signal in each quadrant and compiled the intensity profiles. We observed that the intensity and duration of each calcium pulse correlated with a contraction of the corresponding muscle cell (Fig. 1D). Hence, GCaMP signals represent a good proxy for muscle contractions in the embryo. We subsequently extracted several parameters such as the duration of the calcium pulses (Fig. 1E), the delay between pulses among adjacent cells, (Fig. 1F), and their intensity (Fig. 1G, Table S1). We observed (Movie2), that each pulse lasted roughly 5-6 seconds in cells-1/5 (Fig. 1E). The intensity of contractions differed among cells, with the highest intensity observed in cell-2 and cell-3 (Fig. 1G), which pulsed first in 25 out of 29 cases (in 3/25 cases they pulsed with cells-1/4/5); subsequently, the calcium activity shifted to muscle cells-1/4/5 (Fig.1E; Table S1). Considering the higher pulsing intensity and the fact that they pulsed first, we propose that cell-2 and cell-3 act as leader cells (Fig. 1E-G), as indeed they are different from from cells-1/4/5 (Fig. 1H).

In conclusion, our data reveal the existence of five cells initiating the wave of muscle contractions.

### Cell-2 and cell-3 lead the calcium wave and muscle activity

To clarify the role of cell-2 and cell-3, we ablated these cells by expressing a miniSOG construct (Xu and Chisholm, 2016), which induces photo-damage through the formation of active oxygen species when illuminated at 458 nm. Expressing the miniSOG construct only in muscle cells (Fig. 2A), we specifically photo-ablated cell-2 and cell-3 together in dorsal and ventral left quadrants before muscle contractions started, and analyzed if muscle activity was impaired. As controls, we also photo-ablated three areas of similar size located either in head muscles, tail muscles or head neurons. We reasoned that in kymographs plotting the intensity profile along a line passing through cell-2 and cell-3 in both quadrants (Fig. 2B) a signal along this line should be visible due to embryo rotations (Movie 3). We observed that targeted ablation of cell-2 and cell-3 in two quadrants impaired muscle contractions and embryo rotations (Fig. 2C-D). When similar ablations were made in control areas, muscle activity remained normal (Fig. 2D). In addition, ablation of cells-2/3 resulted in slower elongation (Fig. 2E-F). This result confirms that cell-2 and cell-3 initiate the calcium wave and further suggests that they act as leader cells during muscle contractions.

**Figure 2.**
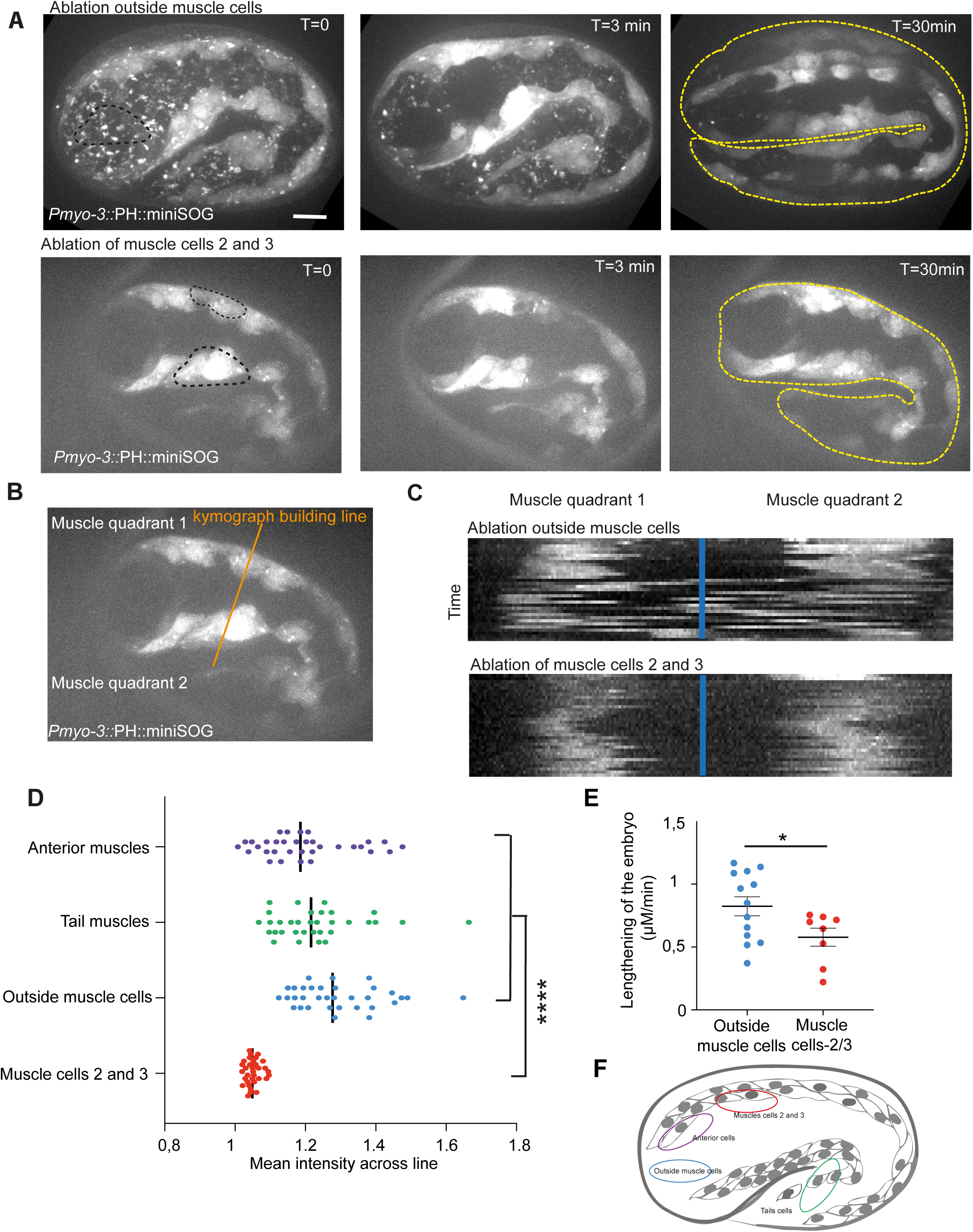
Cell-2 and cell-3 behave as leader cells for muscle activity. **(A)** Snapshot of a timelapse showing a control embryo (top), and an embryo (bottom) in which cells-2/3 were photo-ablated in the dorsal and ventral left quadrants (black dotted lines). Yellow dotted line, embryo contours. **(B)** Localisation of the orange line used for building the kymograph. **(C)** Kymograph for control (top) and ablated cell-2/3 (bottom). The intensity was measured along the blue line. **(D)** Graph presenting the intensity over time. **(E)** Graph presenting the lengthening of the embryo over time (control, n=12 embryos; cell-2/3, n=8 embryos). **(F)** Scheme of an embryo representing the different ROIs used for photoablation (outside, n=12 embryos; cell-2/3, n=12 embryos; anterior muscles, n=7 embryos; tails muscles, n=12 embryos). p-values, *, p<0,05 ****, p<0,0001.

### Two innexins coordinate muscle activity

To determine how muscle contractions are controlled and coordinated among the four quadrants, we performed a gene candidate RNAi screen, focusing on three gene families involved in cell-cell communication: the innexins (components of invertebrate gap junctions), the DEG/ENAC channels some of which are mechanoresponsive ion channels, and TRP channels expressed in embryos, amounting to 47 candidate genes (Table S2). From this screen, we identified two innexins leading to a defect in muscle activity, INX-18 and INX-15. We confirmed their role using the strong mutant *inx-18(ok2454)* (Voelker *et al*., 2019) and a newly CRISPR-induced mutant *inx-15(syb5414)* deleting the first three transmembrane domains (Fig. S2A).

We confirmed that INX-18 is expressed in muscle cells by using a GFP translational (Fig. 3A) and transcriptional reporter constructs (Fig. S2B) (Liu *et al*., 2013). In *inx-18* knockdown and *inx-18(ok2454)* embryos, we observed a ventral bump with more constricted muscle cells (Fig. S2C-D). We further observed that the pulsing delay of cells-2/3 versus cells-1/4/5 was reduced (Fig. 3B), GCaMP intensity was lower (Fig. 3C), and calcium wave was affected (Fig. 3D, Movie4), although the pulse duration was unaffected (Fig. 3I). The reduced distinction between cells-2/3 and cells-1/4/5 (Fig. 3J; compare with Fig. 1H) and the less efficient muscle contractions (Fig. S2H-I) suggest that cells-2/3 have a partially compromised leader cell activity. Moreover, the frequency of contractions was slightly reduced compared to wild-type embryos (Fig. S2F-G).

**Figure 3.**
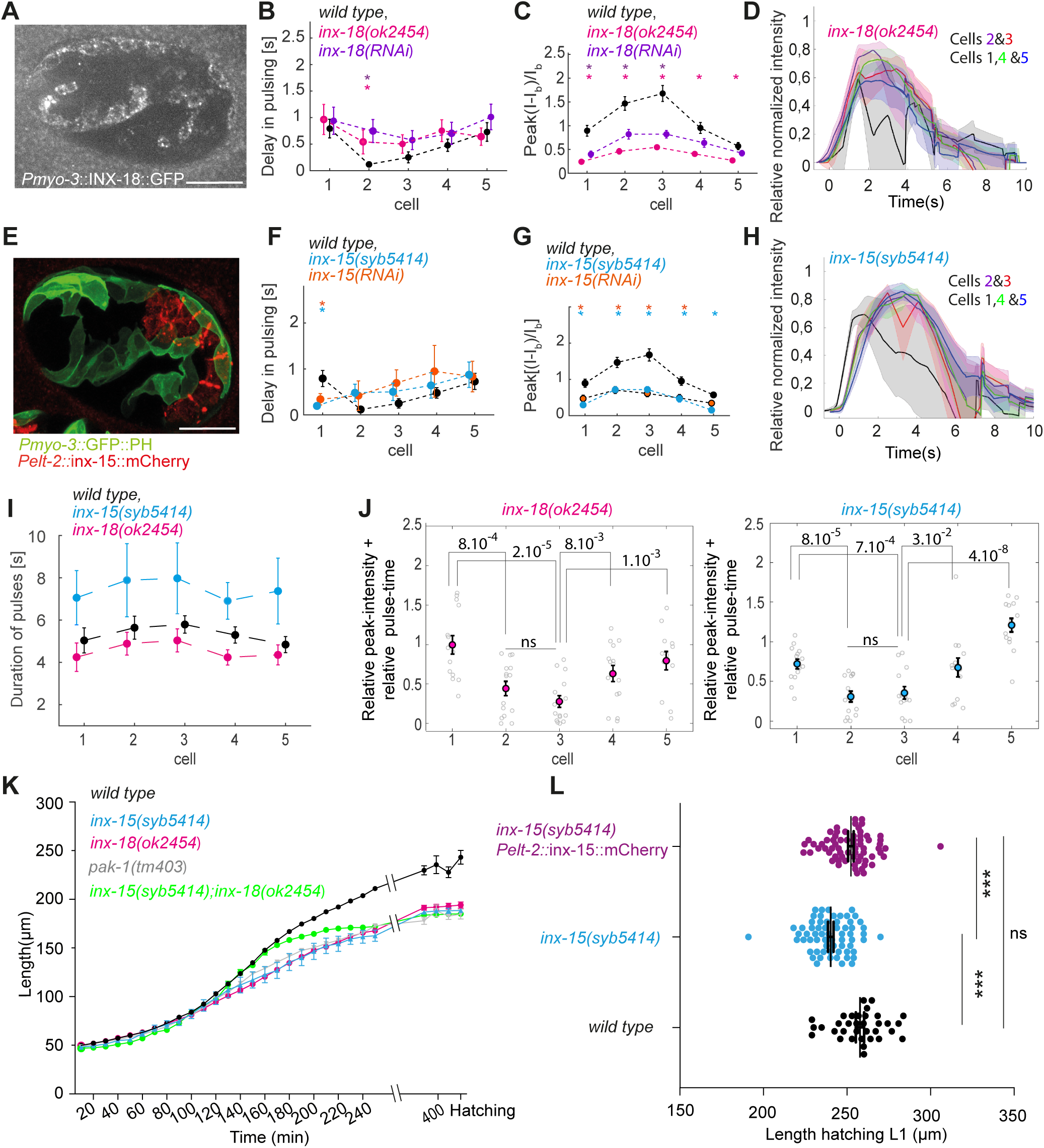
Innexin roles in muscle activity. **(A)** *Pmyo-3::inx-18::*GFP expression in muscles. Note the dotty pattern outlining muscle cells. **(B-C)** Graphs detailing muscle GCaMP in wild type (data from Fig. 1), *inx-18(RNAi)* (n=10 embryos/ 19 waves), and *inx-18(ok2454)* (n=6 embryos/15 waves) embryos, showing delay in pulsing (B) and peak intensity (C). **(D)** Graph showing the normalized calcium wave in *inx-18(ok2454)* embryos. **(E)** Embryo showing *Pelt-2::*INX-15::mCherry and *Pmyo-3*::PH::GFP localization. **(F-G)** Graphs detailing muscle activity in wild type (data from Fig. 1), *inx-15(RNAi)* (n=5 embryos and 12 waves), and *inx- 15(syb5414)* (n=6 embryos and 14 waves) embryos, showing delay in pulsing (F) and peak intensity (G). **(H)** Graph showing the normalized calcium wave in *inx- 15(syb5414*) embryos. **(I)** Plots showing the duration of the pulses in wild type, *inx-18(ok2454)* and *inx-15(syb5414)* embryos. **(J)** Plot showing the relative peak intensity and relative pulse time combined together for *inx-18(ok2454)* and *inx- 15(syb5414)* embryos. **(K)** Elongation curves of wild type, *inx-18(ok2454)*, *inx- 15(syb5414), inx-15(syb5414); inx-18(ok2454)* and *pak-1(tm403)* embryos. **(L)** Length of wildtype, *inx-15(syb5414); Pelt-2*::INX-15::mCherry, and *inx15(syb5414*) hatching L1 larvae (n=32 hatching larvae for all conditions). P- values, ****, p<0,0001.

Interestingly, INX-15 was only expressed in intestinal cells, as previously reported (Altun ZF *et al*.,2009), showing a continuous distribution between adjacent intestinal cells and a punctate one basally (Fig. 3E, Fig. S2E). Analysis of the calcium profile in *inx-15(RNAi)* and *inx-15(syb5414)* embryos revealed, as in *inx- 18* mutants, that the delay, intensity, calcium wave and muscle contractions were affected (Fig. 3F-J, Fig.S2H, Table S1, Movie5). Although cell-1 pulsed first in 5 waves out of 14 analyzed, it did so with a lower intensity, which led us to exclude it as a potential leader cell (Fig. 3J, Table S1).

Since muscles are important for elongation, we next examined if elongation was perturbed in the absence of INX-18 or INX-15 (Fig. 3K). We found that the muscle-dependent phase of elongation of *inx-15(syb5414)* or *inx-18(ok2454)* embryos was slower and that their final length was slightly shorter than in wild-type embryos. Importantly, expressing an INX-15::mCherry construct under the intestine-specific promoter *elt-2* rescued the length of *inx-15(syb5414)* hatching larvae (Fig. 3L). Intriguingly, the *inx-15* and *inx-18* mutant elongation curves resemble that of a *pak-1* mutant (Lardennois *et al*., 2019), raising the possibility that proper coordination of muscle contraction stimulates PAK-1 activity (Fig. 3K).

Since *inx-15* and *inx-18* mutants lead to similar phenotypes, we examined their interactions by constructing an *inx-15; inx-18* double mutant and analyzing their phenotype. We found that the size of double mutant hatching L1 larvae was similar to that of *inx-15* and *inx-18* single mutants (Fig. 3K). Moreover, although there was a slight but significant difference between the double and each single mutant (Tables S1 and S3), the calcium profile of double mutants was slightly reduced but similar (Fig. S2J-K). Hence, we tentatively suggest that INX-15 and INX-18 are acting in the same pathway. Altogether these results highlight a new role of the innexin family in muscle activity control and reveal that the intestine is also required for normal embryonic elongation, much like it also coordinates muscle activity during defecation (Peters *et al*., 2007).

### Two Deg/Enac channels coordinate muscle activity

Although we did not identify any DEG/ENAC channel leading to a muscle phenotype in RNAi screen, the further analysis of two gain of function mutants*, deg-1(u506)* and *unc-105(n506)* (Park and Horvitz, 1986; Liu, Schrank *et al*, 1996; Takagaki *et al*., 2020), uncovered their role in regulating muscle contractions.

Analysis of *deg-1(u506)* showed that cell-1 to cell-5 displayed individual signals instead of strong coordinated pulses as in wild-type embryos (Fig. 4A-C, E, Movie6). The distinction between cells-1/5 was essentially lost and the resulting contraction was weak (Fig. 4F, Fig. S3D). Furthermore, we confirmed that *deg- 1(u506)* embryos arrest at the 2-fold stage at 16°C (Fig. 4G) (García-Añoveros, *et al*, 1995). Our analysis of a DEG-1::GFP knockin showing a sparse dotty pattern (Fig. 4I) colocalizing with muscle membrane markers, near the overlying ECM, (Fig. S3A) further suggests that DEG-1 acts in muscle. We conclude that normal DEG-1 activity is required to coordinate muscle contractions and to ensure normal leader activity, both of which are essential for normal elongation.

**Figure 4.**
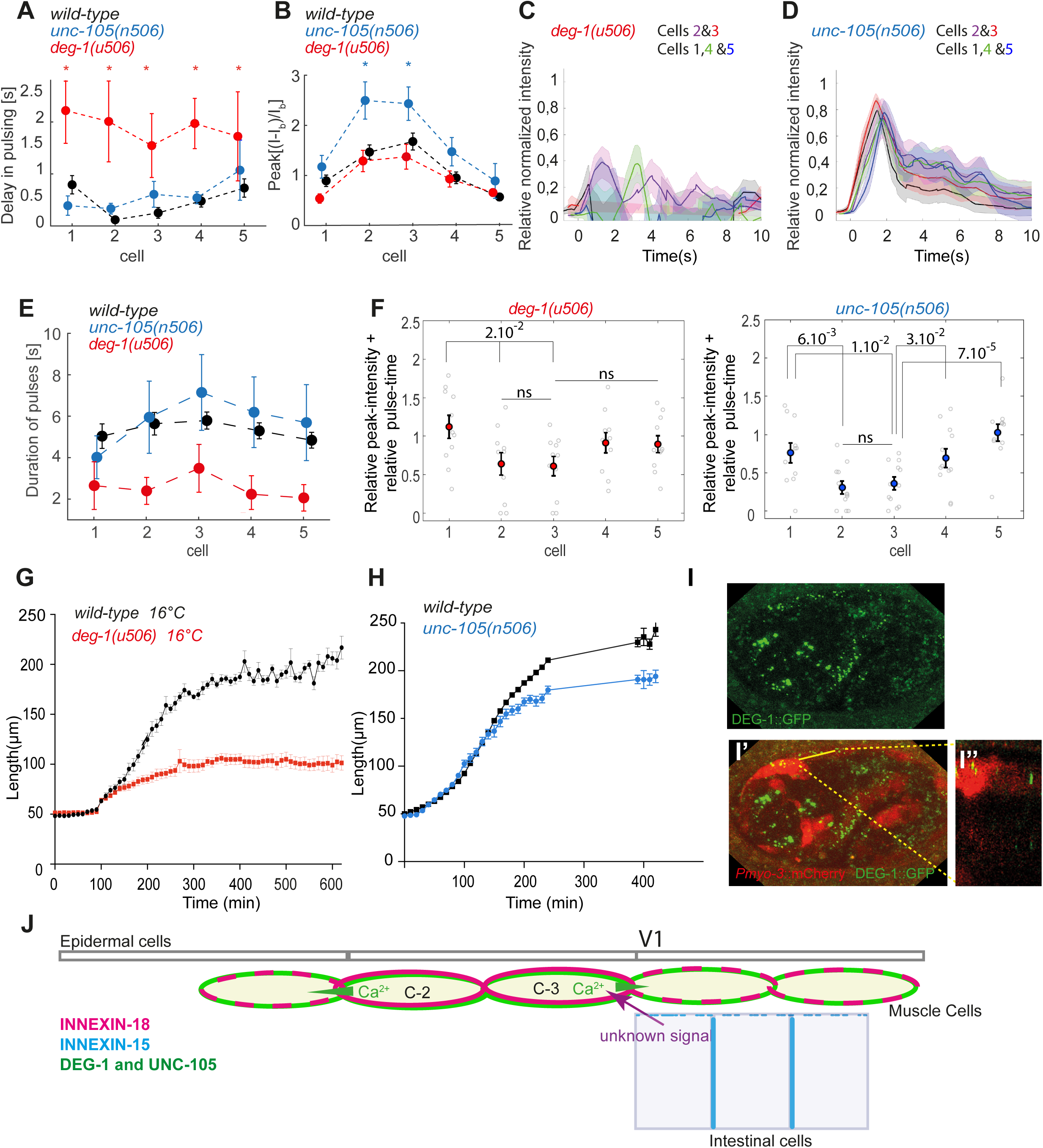
DEG/ENac channels are required for normal muscle activity. **(A-B)** GCaMP activity analysis showing the delay in pulsing (A), the peak intensity of the pulses (B) for wild-type (data from Fig. 1), *unc-105(n506)* (n=8 embryos and 10 waves) and *deg-1(u506)* (n=8 embryos, 12 waves.) **(C-D)** Graph showing the normalized calcium wave in *deg-1(u506)* (C) and *unc-105(n506*) (D) embryos. **(E)** Plots showing the duration of the pulses in wild type in wild-type, *deg-1(u506)* and *unc-105(n506).* **(F)** Plot showing the normalized intensity and delay combined together *deg-1(u506)* and *unc-105(n506)* embryos. **(G)** Elongation curve for *deg-1(n506)* (n=9) and wild-type(n=10) at 16°C**. (H)** Elongation curve for *unc-105(n506)* (n=6) and wild-type (n=12) at 20°C. Data in (G-H) represent mean values ± SEM. **(I)** Localisation of DEG-1::GFP in muscle cells. **(I’)** Merged images of DEG-1::GFP and *Pmyo-3::*mCherry , **(I”)** Transverse view of the image in (I’). **(J)** A model for the coordination of muscle activity in embryos. The proposed model involves an unknown signal implicating INX-15 in intestinal cells, which preferentially activates muscle cells-2 and -3 through INX-18. Of note, how muscle cells initiate their activity remains unknown. INX-18 facilitates signal propagation and activates surrounding muscle cells. Within the muscle quadrant, the activities of DEG-1 and UNC-105 are essential for maintaining synchronization between muscle cells and aiding signal propagation. Together, these interactions among the three tissues ensure proper embryo elongation at the level of epidermal cells.

We found that the *unc-105(n506)* gain-of-function mutation resulted in embryos with normal contraction time and delay in pulsing (Fig. 4A-E, Movie7). However, a higher calcium intensity peak was observed in cell-2 and cell-3 compared to wild-type embryos (Fig. 4B). In addition, relaxation seemed to be impaired, as both the Ca^++^ (Fig. 4D) and contraction (Fig. S3C) profiles did not return to baseline as quickly as in wild-type, resulting in slightly blunted difference among cells-1/5 (Fig. 4F). Consistently, the integral calcium intensity computed on the wave duration was the highest among genotypes (Fig. S3D). Furthermore, embryonic elongation was affected resulting in shorter terminal embryos than controls (Fig. 4H), presumably due to the relaxation defects and increased contraction frequency (Fig. S2F). The analysis of an *unc-105::GFP* knockin strain showed a widely distributed expression, which did not appear to be exclusively located in muscle cells (Fig. S3B). Overall, these results reveal a novel role for UNC-105 during embryonic elongation, which is important for relaxing muscles after contraction.

### A model for the coordination of muscle contraction

Taken together, our results show that intestinal cells are involved in the morphogenesis of the embryo during the second stage of elongation. This new role for intestinal cells in the overall elongation of the embryo highlights the requirement for cells of the three germ layers in ensuring proper morphogenesis. The nature of the signal initiating muscle contractions and the molecular mechanism by which intestinal cells control muscle activity will await further studies. Indeed, we could not identify any calcium exchange between intestinal and muscle cells, nor a clear apposition between the two innexins even if they are close (Fig. 3E), nor identify their origin. One possibility is that ions other than Ca^++^ are involved, such as H^+^ ions as in defecation (Pfeiffer *et al*., 2008; Wagner *et al*., 2011), or that the probes used in our attempts were not sensitive enough. Previous work on *elt-2* mutants, which affect intestine differentiation, did not report an elongation defect (Fukushige *et a.l*, 1998), yet a small size reduction as observed in *inx-15* mutants could have been missed.

The simplest model to explain the reduced leadership of cell-2 and cell-3 in *inx- 18* and *inx-15* mutants is that a signal emitted by intestinal cells through INX-15 singles out cells-2/3 (Fig. 4J), which in turn activate cells-1/4/5. This model accounts for the fact that *inx-15* and *inx-18* may act in the same pathway and for the respective localization of both proteins. In addition, INX-18 could be involved in communication between muscle cells. The signal transmitted by INX-15 and INX-18, and its propagation, subsequently require the Deg/Enac channels DEG-1 and UNC-105 to synchronize muscle cell activity in one quadrant. The effect on embryonic elongation observed in all four mutants suggests that the duration and magnitude of muscle contractions, along with muscle relaxation, are likely required to generate a proper mechanotransduction response in the epidermis. As DEG-1 is located at the interface between muscle cells and the surrounding collagen matrix, we could hypothesize that its opening in response to mechanical input balances and retro-controls muscle activity to allow muscle cell coordination dynamics (Liu *et al.,* 1996; Yan *et al*., 2020).

In summary, this analysis highlights the role of adjacent cell bioelectric coupling, involving gap junctions and Deg/Enac channels, in facilitating cooperation between different connected tissues.

## Materials and methods

### Strains

Animals were grown on NGM (nematode growth medium) plates with a bacterial lawn of the OP50 E. coli strain, following the protocol of Brenner (1974) and generally maintained at 20°C unless otherwise stated for temperature sensitive mutants. The alleles *deg-1(syb5346)*, strain PHX5346, *inx-15(syb5728)*, strain PHX5728 and *unc-105(syb4276)*, strain PHX4276, as well as the knockin *deg-1::GFP(syb4251)*, strain PHX4251, and were generated by Suny Biotech (Fu Jian Province, China; https://www.sunybiotech.com/) using the CRISPR/CAS9 technology (table S4). The allele *inx-15(syb5728)*, strain PHX5728, is a deletion of 904 bp (first two exons and first intron) corresponding to AA 11 to AA 220 deletion of the 3 transmembrane domains.

### RNAi mini-screen

RNAi was performed by feeding L4 hermaphrodites with HT115 E. coli expressing dsRNA. 41 dsRNA cloned in the L440 vectors came from the Ahringer’s library were sequenced to confirm the target gene. The other dsRNAs were cloned into the L440 vector after amplification of the sequence using the appropriate set of primers and subsequent transformation in HTT115 E*. coli*. L4 animals were fed during 24 h or 48 h hours prior to the analysis the progeny.

### Real-time calcium intensity analysis

The embryos were mounted on a glass slide over a 2% agarose pad with the M9 buffer (86 mM NaCl, 42 mM Na2HPO4, 22 mM KH2PO4, 1 mM MgSO4).

Embryos were observed with a Zeiss Axio Observer.Z1 inverted spinning disk microscope equipped with a *CSUX1-A1* (Yokogawa) spinning disc module, a Roper Evolve camera and an apochromatic oil-immersion 100x objective (NA=1.4). The microscope was controlled by the Metamorph (Molecular Devices) acquisition software. For long-time imaging of *Pmyo3*::PH::GFP membrane labeled muscle, excitation was set at 100 ms and 20 % of laser at 2 min intervals. For calcium imaging, a stream acquisition mode was used with 50 ms of excitation and 36,5 % of laser at 89 ms intervals. All live imaging experiments were performed at 20°C, except for experiments involving *deg- 1(u506)* which were performed at 16°C.

### Photoablation experiment

To specifically ablate cells 2 and 3, we used a transgenic animal carrying the *Pmyo3*::MiniSOG::mCherry construct which is active only in muscle cells. Just prior to the start of muscle activity, we designed an ROI (region of interest) for cells 2 and 3 in the ventral and dorsal quadrants, took two images 1 min and 30 s before photoablation. We then photo-ablated the two cells in each quadrants using the FRAP module of the ILAS software installed on the spinning disc with a 456 nm laser at 80% power for 2 min, and finally recorded muscle activity every 30 s for 25 minutes. For the control experiment, we ablated an ROI located outside the *myo-3* labelled cells, but in the same focal plane as muscles; this ROI devoid of any labelling could correspond to the location of the neurons.

### Photo Ablation analysis

To quantify muscle activity and muscle twitching, we used the Image J software (NIH, Bethesda, Maryland, USA; http://rsb.info.nih.gov/ij/) to design a line passing through cells 2 and 3 and built a kymograph.

### Analysis of elongation rate

DIC time-lapse videos were obtained using a Leica DM6000 microscope controlled by a Leica LAS-AF software. Observations were done under a 40X∼ oil immersion objective. Mothers were cut up to gain early-staged embryos, which were then transferred onto thin 5% soft agarose pads in a drop of M9. Z-stack image series with a 1.5-µm Z-step distance were taken every 10 min during 16 h. The image J software was used to quantify the embryonic length from the end of ventral enclosure/onset of elongation, by taking a segmented line through the midline of the embryos from head to tail. For the *deg-1(u506)* mutant we set up the microscopy room at 16°C.

### Fluorescence microscopy analysis

We acquired fluorescence images with a LSM 980 Airyscan2 Zeiss confocal system equipped with 488 nm and 561 nm excitation laser lines and a 63x oil immersion objective with NA 1,4 and 0,15 µm z-steps. In confocal mode, we performed a simultaneous acquisition for the green and red channels in order to get both markers at the same time. These images were deconvoluted with the Huygens software, then analyzed single plane images or max z projection images using the Image J software.

### Calcium image analysis

We performed maximal projection along z of the original time-dependent movies of HBR4 and corrected them for photobleaching by using the dedicated exponential correction tool in FIJI. Subsequently, we used a Matlab script to measure the average fluorescence intensity in cell cytoplasm by hand-tracking the cells contours by using the *roipoly* function (ROI, region of interest). In each embryo we monitored and tracked the average fluorescence of five muscle’s cells below the H1, H2, v1 cells. Moreover, we recorded the spatial coordinates of the center of mass of each ROI; subsequently we converted them in curvilinear abscissa with respect to the anterior-posterior (AP) axis after manually tracing the body midline from the tip of the embryo head to the tip of the tail. To isolate pulses of signal in a selected cell, we plotted the behavior of the average fluorescence in time. We observed that the signal intensity increased with respect to its baseline within a few seconds, reached a maximum and then decreased again to a value similar to the initial baseline. The resulting time behavior was similar to a bell shape. Indeed, we observed that a pulse in a cell was coordinated with pulses in neighboring cells, which tend to pulse in a group producing a wave of calcium signal. We decided to characterize a single pulse by its initiation time, its termination time and Its duration. The initiation time was calculated as the time at which the calcium pulse intensity reaches 30% of its maximum, as illustrated in Fig. 1D. It was calculated on the intensity normalized between 0 and 1 by subtracting the initial intensity I_0_ which is the initial fluorescence intensity of each cell from I(t) and dividing by the maximum of I(t)-I_0_. The termination time was defined as the time at which the normalized intensity decreased till 30% of its maximum after reaching its maximum. The choice of the threshold of 30% is arbitrary. The duration of a calcium pulse is calculated as the difference between the termination time and the initiation time of the normalized intensity.

To characterize the coordination of the pulses between neighboring cells we measured the delay in pulsing between different cells and the speed of the calcium wave. We estimated the delay with respect to the cell among the five cells pulsing first. More precisely the delay in pulse was defined as the difference in the initiation time with the first pulsing cell. We measured the average wave velocity as *v_w_= abs(<x_last_>-<x_first_>)/(<t_last_>-<t_first_>)*, where *(<t_last_>-<t_first_>)* is the difference between the average delay at which the first cell (*<t_first_>*) pulsed and the average delay at which pulse the last cell (*<t_last_>*); *abs(<x_last_>-<x_first_>)* is the modulus of the difference between the average AP position of the last pulsing cell *<x_last_>* and the average AP position of the first pulsing cell *<x_first_>*. Finally, we characterized the intensity of the pulses by measuring the peak intensity as the maximal intensity reached by a cell during a pulse normalized with respect to the minimal baseline value of the intensity *I_b_* measured among the selected five cells. More precisely we defined *I_peak_* = (*I_max_*- *I_b_*)/ - *I_b_*. Peak [(I-I_b_)/I_b_] where I_b_ is the baseline fluorescence intensity that is chosen as the lowest value of the intensity among the 5 cells in one wave.

To collectively describe the pulse delay and peak intensity so as to capture a potential stereotypical behavior among the first five cells, we added the two variables after normalization as follows.

- The normalised relative peak intensity was *abs(I_peak_-max(I_peak_))/max(I_peak_)*, with *I_peak_ =Peak[(I - I_b_)]/I_b_]*; *abs,* absolute value.
- The normalised relative pulsing time was *(T_p_ -min(T_p_))/max(T_p_)*, with *T_p_* the initiation time for the pulse.

Both variables run in the range [0,1) by construction and both are dimensionless; moreover, they have a zero value in a given cell if the cell pulses first or if the cell shows the maximal peak intensity. The results of such plots are presented in Figs. 1H, 3I, 4E.

We defined the integral normalized intensity as the temporal integral of *I*=I_n_-I_n_(t=0)*, where *I_n_=(I - I_b_)]/I_b_*. Practically we summed all the values of *I\** over the first 15 seconds (within this time the majority of pulses takes place). This variable captures the intensity of pluses and their duration at the same time.

All the MATLAB code are available upon request

### Statistical analysis

For elongation curves, the standard error of the mean (SEM) was calculated. For L1 hatching measurements and ablation intensity, we performed paired t-tests or ANOVA test using the GraphPad PRISM software. For calcium analysis, all the analyses were done using MATLAB (The MathWorks Inc., Natick, MA). All the p values are in table S3.

## Acknowledgements

The authors thank Jean-Louis Bessereau, and an anonymous reviewer for their careful reading of the manuscript, the Caenorhabditis Genetics Center (funded by the NIH Office of Research Infrastructure Programs P40 OD010440), Sunybiotech, Andrew D. Chisholm, Henrik Bringmann, Jihong Bai, Shinji Ihara Jihong Bai for strains and reagents, and the IBPS imaging center for advice. This work was supported by grants from the Agence Nationale pour la Recherche (grant #ANR-18-CE13-0008-01) and the Association pour la Recherche sur le Cancer (Subvention 2018).

## Authors’ contributions

FL and ML conceived the project and wrote the manuscript. FL performed most experiments; TF performed Ca++ quantifications, XY quantified embryo rotations and HL the connections between muscle quadrants.

## Supplementary Figure legends

**Supplementary Figure 1. Characterization of the first contracting muscles. (A)** Snapshots from a *Pmyo-3*::GCaMP::GFP timelapse showing the first muscle contraction which occurs dorsally activity (see Movie1). **(B)** Fluorescent embryo showing the respective localization of cell-2 and cell-3 (*Pmyo-3*::GCaMP::GFP) compared to pharynx PHA-4::GFP positive cells(arrow). **(C)** Fluorescent embryo showing the respective localization of cell-2 and cell-3 (*Pmyo-3*::GCaMP::GFP) compared to epidermal DLG-1::RFP positive cells.

**Supplementary Figure 2. Additional characterization of INX-15 and INX-18. (A)** Position of *inx-15(syb5414)* deletion. **(B)** Distribution of a *Pinx-18::gfp* translational reporter. **(C-D)** Embryos labelled with the *Pmyo-3*::pH::GFP in control (C) *and inx-18(RNAi)* background. White arrow, bump due to ventral constriction (D). **(E)** Embryo showing the *Pinx-15::GFP* transcriptional reporter profile. **(F)** Contraction frequency (number of contractions/minute) in wild-type, *deg-1(u506)*, *unc-105(n506), inx15(syb5414)*, and *inx-18(ok2454)*. **(G)** Propagation speed of the calcium wave in *wild type*, *unc-105(n506), inx15(syb5414)*, and *inx-18(ok2454)* embryos. **(H)** Graphs showing the time correlation between muscle contraction (distance between cell-1 and cell-3) in blue and the normalized GCaMP intensity of cell-3 in red in *inx15(syb5414)*, and *inx-18(ok2454)* embryos. **(I)** Plots showing the temporal integral of the normalized intensity in wild type, *inx-18(ok2454)* and *inx-15(syb5414)* (see methods). **(J-K)** Graphs showing the delay in GCaMP signal (J), and the peak intensity of the pulses (K) for wild-type, *inx-18(ok2454)*, *inx-15(syb5414)* and *inx- 15(syb5414); inx-18(ok2454)* embryos. Scale bar, 10µm.

**Supplementary Figure 3. Additional characterization of DEG-1 and UNC-105. (A)** Images of DEG-1::GFP (green), EMB-9::mCherry (red), and merge. **(B)** images of UNC-105::GFP (green), EMB-9::mCherry (red), and merge. Scale bar, 10 µm; yellow arrow, muscle localization **(C)** Graphs showing the time correlation between muscle contraction (distance between cell-1 and cell-3) in blue and the normalized GCaMP intensity of cell-3 in red in wild type, *deg- 1(u506)* and *unc-105(n506).* T*, time of minimal distance between cell-1 and - 3.

**Supplementary Movie 1**: Movie of a pre-contraction embryo expressing *Pmyo- 3*::GCaMP::GFP; initially the GCaMP signal is randomly blinking in all cells, then a strong signal in five dorsal cells accompanying the first contraction. Scale bar, 10 µm.

**Supplementary Movie 2**: Movie of a contracting embryo expressing more strongly *Pmyo-3*::GCaMP::GFP in five cells within different muscle quadrants. Scale bar, 10 µm.

**Supplementary Movie 3**: Movie of a control embryo (left), and an embryo (right) in which cell-2 and cell-3 were photoablated in the dorsal and ventral left quadrants expressing the *Pmyo-3*::PH::miniSOG. Note how the control embryo, but not the ablated embryo, rotates within the eggshell. Scale bar, 10 µm.

**Supplementary Movie 4**: Movie of an embryo expressing *Pmyo-3*::GCaMP::GFP in *inx-18(ok2454)* embryo. Note that the signal is weaker. Scale bar, 10 µm.

**Supplementary Movie 5**: Movie of an embryo expressing *Pmyo-3*::GCaMP::GFP in *inx-15(syb5414)* embryo. Scale bar, 10 µm.

**Supplementary Movie 6**: Movie of an embryo expressing *Pmyo-3*::GCaMP::GFP in *deg-1(u506)* embryo. Note the rapid succession of GCaMP signals in different places, and the absence of rotation. Scale bar, 10 µm.

**Supplementary Movie 7**: Movie of an embryo expressing *Pmyo-3*::GCaMP::GFP in *unc-105(n506)* embryo. Scale bar, 10 µm.

