## Supplemental 3 figures and 4 tables for "Muscle and intestine innexins with muscle Deg/Enac channels promote muscle coordination and embryo elongation": Llense-etal_supp.pdf

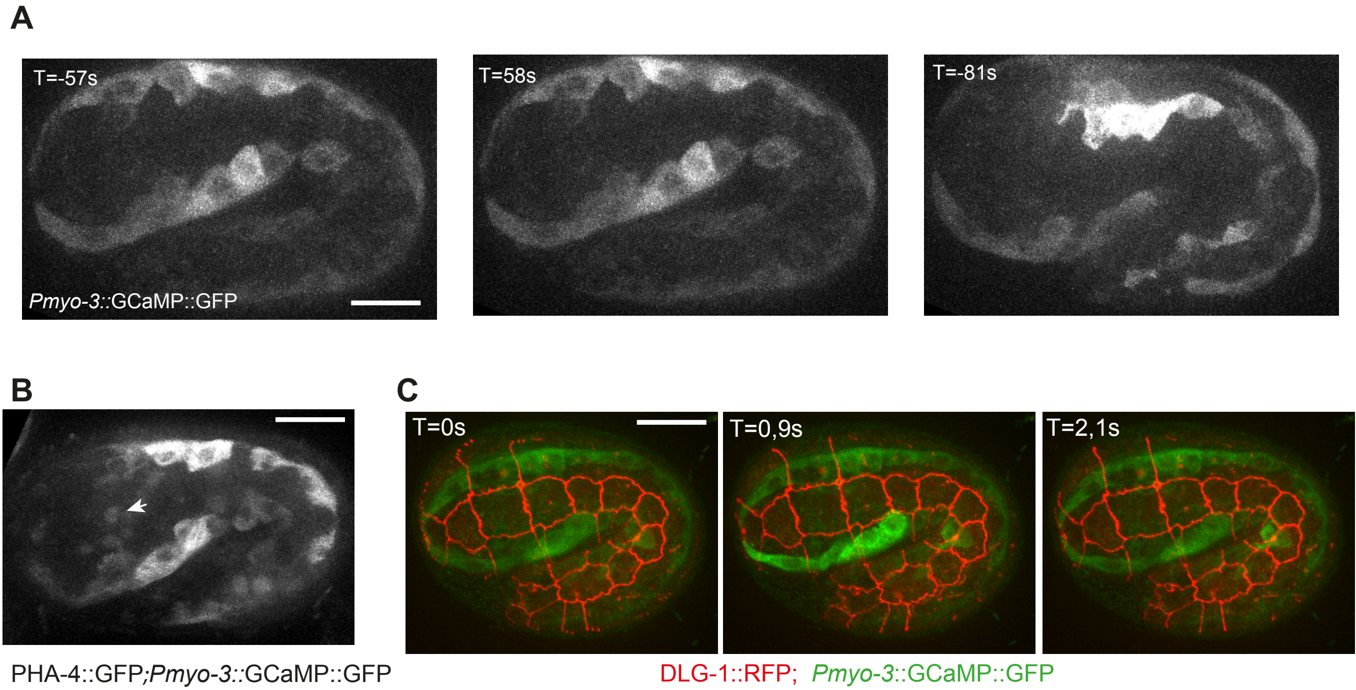
A
T=-57s
Pmyo-3::GCaMP::GFP
T=58s
T=-81s
B
PHA-4::GFP;Pmyo-3::GCaMP::GFP
C
T=0s
T=0,9s
T=2,1s
DLG-1::RFP; Pmyo-3::GCaMP::GFP

Figure S1

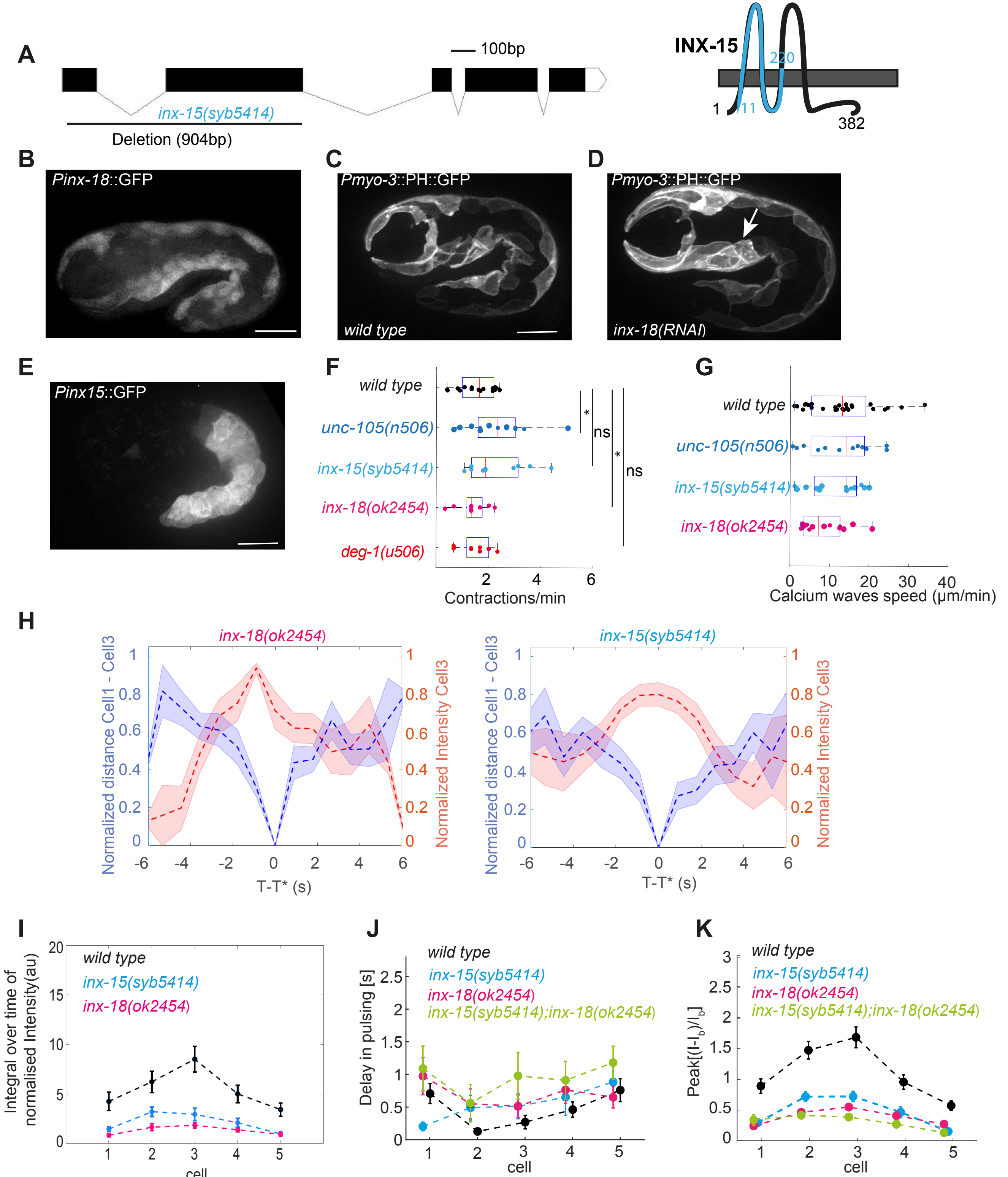

A
100bp
inx-15(syb5414)
Deletion (904bp)
INX-15
220
1
111
382
B
Pinx-18::GFP
C
Pmyo-3::PH::GFP
wild type
D
Pmyo-3::PH::GFP
inx-18(RNAi)
E
Pinx15::GFP
F
wild type
unc-105(n506)
inx-15(syb5414)
inx-18(ok2454)
deg-1(u506)
ns
ns
Contractions/min
G
wild type
unc-105(n506)
inx-15(syb5414)
inx-18(ok2454)
Calcium waves speed (µm/min)
H
inx-18(ok2454)
inx-15(syb5414)
Normalized distance Cell1 - Cell3
Normalized Intensity Cell3
T-T* (s)
I
Integral over time of normalised Intensity(au)
wild type
inx-15(syb5414)
inx-18(ok2454)
cell
J
Delay in pulsing [s]
wild type
inx-15(syb5414)
inx-18(ok2454)
inx-15(syb5414);inx-18(ok2454)
cell
K
Peak[(I-I_b)/I_b]
wild type
inx-15(syb5414)
inx-18(ok2454)
inx-15(syb5414);inx-18(ok2454)
cell


Figure S2

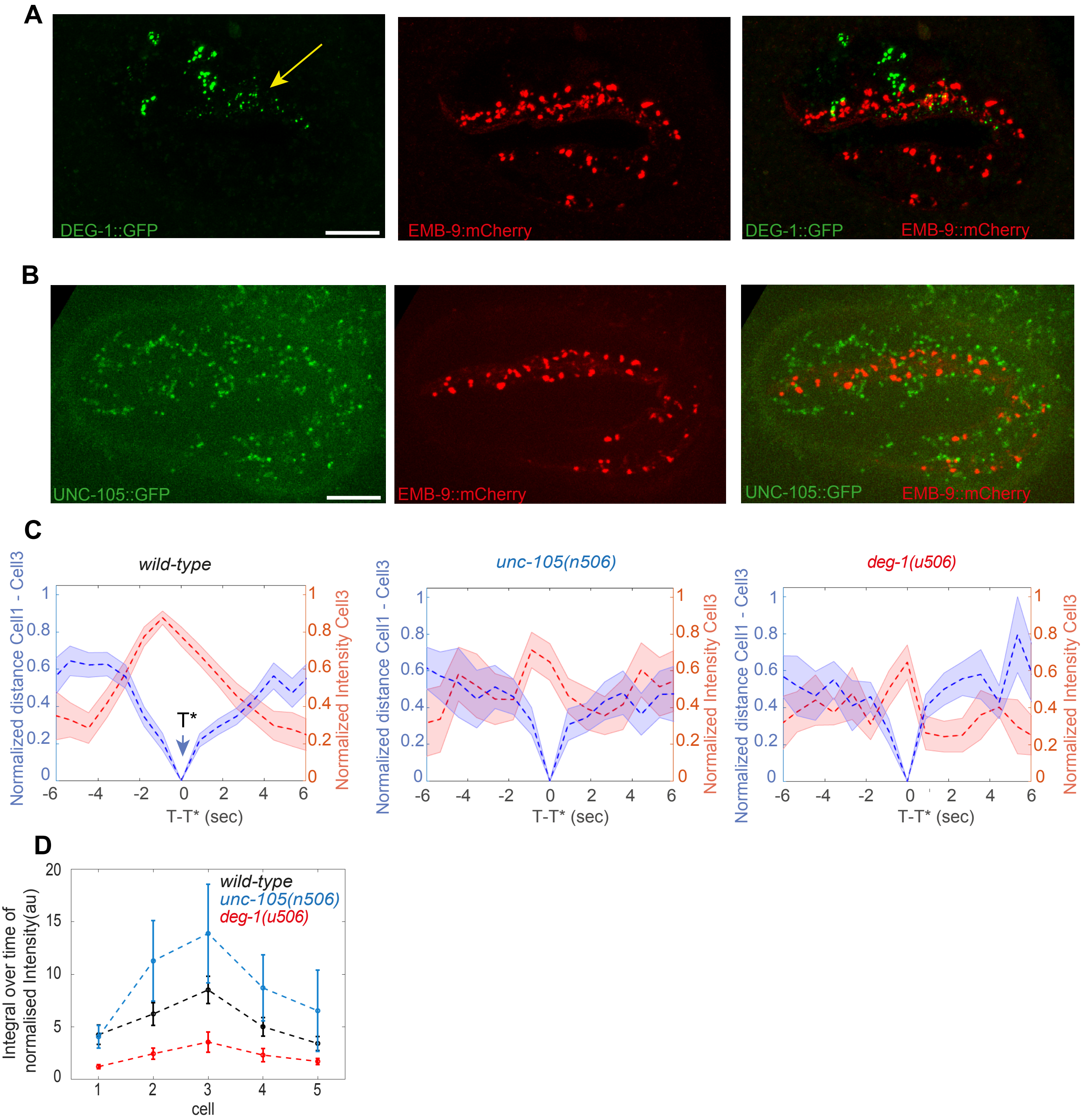
A
DEG-1::GFP
EMB-9:mCherry
DEG-1::GFP
EMB-9::mCherry
B
UNC-105::GFP
EMB-9::mCherry
UNC-105::GFP
EMB-9::mCherry
C
wild-type
unc-105(n506)
deg-1(u506)
Normalized distance Cell1 - Cell3
Normalized Intensity Cell3
T*
T-T* (sec)
D
wild-type
unc-105(n506)
deg-1(u506)
Integral over time of normalised Intensity(au)
cell

FigureS3

**Table S1**: Delay and intensity for all genotypes

*1-wild-type* (13 embryos, 29 waves)

| Delay | | | | | | Intensity | | | | |
|---|---|---|---|---|---|---|---|---|---|---|
| Cell1 | Cell2 | Cell3 | Cell4 | Cell5 | | Cell1 | Cell2 | Cell3 | Cell4 | Cell5 |
| 0.1 | 0.5 | 0 | 0 | 1 | **3**=4-1-2-5 | 1.75 | 1.60 | **2.51** | 0.77 | 0.66 |
| 0 | 0.05 | 0.1 | 0.6 | 1.7 | **1**-2-3-4-5 | **1.66** | 1.36 | 1.51 | 0.44 | 0.36 |
| 0 | 0.2 | 0.15 | 0.1 | 1.05 | 1-4-**3**-2-5 | 1.79 | 1.41 | 1.85 | 0.62 | 0.53 |
| 0.7 | 0 | 0 | 0.9 | 0.75 | 2=**3**-1-5-4 | 2.07 | 2.35 | 4.07 | 0.76 | 0.74 |
| 1.35 | 0.45 | 0.65 | 0.3 | 0 | 5-4-2-**3**-1 | 0.39 | 0.41 | 1.72 | 0.94 | 0.51 |
| 0.1 | 0.05 | 0 | 0.05 | 0.05 | 3-**2**=4=5-1 | 2.23 | 3.32 | **3.10** | 1.24 | 1.91 |
| 0.45 | 0 | 0.4 | 1.05 | 0.55 | **2**-3-1-5-4 | 0.89 | **1.96** | 1.67 | 0.98 | 0.90 |
| 0.15 | 0 | 0.1 | 0.3 | 0.15 | **2**-3-1-4-5 | 0.38 | **1.82** | 1.80 | 0.98 | 0.89 |
| 0.3 | 0.1 | 0 | 0.6 | 0.45 | 3-**2**-1-5-4 | 0.99 | 1.96 | **1.76** | 1.01 | 0.85 |
| 0.65 | 0 | 0.1 | 0.45 | 1.3 | **2**-3-4-1-5 | 0.30 | **0.61** | 0.61 | 0.43 | 0.18 |
| 3.45 | 0.15 | 0 | 0.75 | 0.2 | 3-**2**-5-4-1 | 1.08 | 1.54 | 0.94 | 0.60 | 0.29 |
| 0.9 | 0 | 0.85 | 0.9 | 1.35 | 2-**3**-1-4-5 | 0.58 | 0.74 | 0.91 | 0.55 | 0.22 |
| 1.3 | 0.15 | 0 | 0.75 | 0.2 | 3-**2**-5-4-1 | 1.08 | 1.54 | 0.94 | 0.60 | 0.29 |
| 0.4 | 0 | 0.65 | 1.15 | 3.85 | **2**-1-3-4-5 | 0.07 | 0.77 | 0.59 | 0.16 | 0.01 |
| 0.9 | 0.25 | 0 | 0.05 | 0.05 | **3**-4=5-2-1 | 0.63 | 1.15 | 1.33 | 1.26 | 0.59 |
| 0.45 | 0 | 0 | 0.1 | 0.05 | **2**=3-5-4-1 | 0.40 | 1.60 | 1.37 | 1.04 | 0.68 |
| 0 | 0.1 | 0 | 0.5 | 0.45 | 3=1-**2**-5-4 | 0.93 | 3.29 | **2.92** | 0.40 | 0.12 |
| 0.2 | 0 | 0.05 | 0.2 | 0.5 | 2-**3**-4=1-5 | 0.87 | 2.13 | 2.25 | 1.74 | 0.84 |
| 0.3 | 0 | 0.05 | 0.1 | 0.25 | 2-**3**-4-5-1 | 0.78 | 1.99 | 2.23 | 1.66 | 0.82 |
| 0.7 | 0 | 0.05 | 0 | 0.2 | 2-4-**3**-5-1 | 0.96 | 1.95 | 2.17 | 1.72 | 0.86 |
| 0 | 0.05 | 0 | 0.05 | 0.05 | **3**=1-2=4=5 | 1.06 | 1.81 | **2.20** | 1.94 | 0.63 |
| 0.8 | 0.1 | 0.05 | 0 | 0.3 | 4-**3**-2-5-1 | 0.40 | 1.39 | 1.76 | 1.08 | 0.29 |
| 1.85 | 1.25 | 0 | 0.1 | 0.3 | **3**-4-5-2-1 | 0.23 | 0.93 | 1.36 | 0.70 | 0.26 |
| 1.35 | 0 | 0.15 | 0.1 | 0.15 | 2-4-**3**=5-1 | 0.27 | 0.81 | 1.07 | 0.63 | 0.24 |
| 0.45 | 0 | 2.75 | 3 | 1.9 | **2**-1-5-3-4 | 0.96 | 1.10 | 0.22 | 0.50 | 0.65 |
| 0.3 | 0.05 | 0 | 0.05 | 0.1 | 3-**2**=4-5-1 | 0.47 | 1.04 | 0.91 | 0.45 | 0.19 |
| 0.1 | 0 | 0.05 | 0.7 | 0.15 | **2**-3-1-5-4 | 0.65 | 1.26 | 0.96 | 0.49 | 0.18 |
| 2.4 | 0 | 0.8 | 0.95 | 1 | 2-3-**4**-5-1 | 0.11 | 0.14 | 0.54 | 0.58 | 0.38 |
| 3.35 | 0 | 0.35 | 0.1 | 3.1 | 2-4-**3**-5-1 | 1.78 | 1.39 | 3.09 | 3.12 | 1.29 |

Red = min(delay) in cell 1 or 4 or 5

Green = min(delay) in cell 2 or 3

➔ Waves where cells 1 or 4 or 5 pulsed first before cells 2 or 3

4 waves out of 29 (14%) had a first pulsing cell different from cells 2 or 3, and among them only one pulsed with the highest intensity

*2-inx-15(syb5414)* (6 embryos; 14 waves)

| **Delay** | | | | | | **Intensity** | | | | |
|---|---|---|---|---|---|---|---|---|---|---|
| Cell1 | Cell2 | Cell3 | Cell4 | Cell5 | | Cell1 | Cell2 | Cell3 | Cell4 | Cell5 |
| 0.5 | 0.35 | 0 | 0.15 | 0.75 | 3-4-2-1-5 | 0.45 | 0.68 | 1.14 | 0.44 | 0.15 |
| 0 | 0.1 | 1.25 | 0.05 | 0.2 | 1-4-2-3-5 | 0.46 | 0.52 | 1.06 | 0.35 | 0.03 |
| 0.1 | 0.05 | 0.15 | 0 | 0.3 | 4-2-1-3-5 | 0.51 | 0.49 | 0.98 | 0.34 | 0.06 |
| 0.6 | 1.85 | 1.7 | 0 | 0.1 | 4-5-1-2-3 | 0.16 | 0.53 | 0.46 | 0.23 | 0.07 |
| 0.45 | 0 | 0.25 | 0.1 | 0.7 | 2-4-3-1-5 | 0.33 | 1.28 | 1.00 | 1.07 | 0.28 |
| 0 | 0 | 0 | 0.05 | 0.2 | 1=2=3-4-5 | 0.20 | 1.11 | 0.82 | 0.93 | 0.20 |
| 0.25 | 0.05 | 0 | 0.1 | 0.9 | 3-2-4-1-5 | 0.23 | 0.98 | 0.75 | 0.86 | 0.19 |
| 0 | 0.1 | 0.05 | 0.05 | 1.1 | 1-3=4-2-5 | 0.19 | 0.90 | 0.70 | 0.79 | 0.20 |
| 0 | 0.55 | 0.5 | 1.25 | 0.4 | 1-5-3-2-4 | 0.1516 | 0.39 | 0.37 | 0.26 | 0.28 |
| 0.2 | 0.2 | 0 | 0.25 | 0.85 | 3-1=2-4-5 | 0.37 | 0.85 | 0.89 | 0.25 | 0.11 |
| 0 | 2 | 0.1 | 0.6 | 0.35 | 1-3-2-4-5 | 0.34 | 0.76 | 0.77 | 0.17 | 0.08 |
| 0 | 1 | 1.25 | 1.65 | 3.4 | 1-2-3-4-5 | 0.35 | 0.41 | 0.48 | 0.33 | 0.20 |
| 0.05 | 0 | 1.8 | 1.2 | 2.8 | 2-1-4-3=5 | 0.29 | 0.36 | 0.49 | 0.35 | 0.20 |
| 0.6 | 0.5 | 0 | 3.55 | 0.2 | 3-2-5-1-4 | 0.04 | 0.69 | 0.09 | 0.05 | 0.01 |

3- *inx-18(ok2454)* (6 embryos; 15 waves)

| **Delay** | | | | | | **Intensity** | | | | |
|---|---|---|---|---|---|---|---|---|---|---|
| Cell1 | Cell2 | Cell3 | Cell4 | Cell5 | | Cell1 | Cell2 | Cell3 | Cell4 | Cell5 |
| 0.65 | 0 | 0.05 | 0.25 | 0.25 | 2-3_4=5-1 | 0.46 | 0.46 | 0.57 | 0.2 | 0.2 |
| 0.5 | 0 | 0.2 | 0.25 | 1.1 | 2-3-4-1-5 | 0.10 | 0.80 | 0.68 | 0.43 | 0.18 |
| 0.1 | 0.15 | 0 | 0.7 | 1.15 | 3-1-2-4-5 | 0.30 | 0.32 | 0.67 | 0.48 | 0.20 |
| 1.1 | 1.5 | 0 | 1.55 | 0.95 | 3-5-1-2-4 | 0.36 | 0.60 | 0.55 | 0.44 | 0.44 |
| 1.25 | 0.15 | 0 | 1.25 | 1.15 | 3-2-5-1=4 | 0.26 | 0.55 | 0.48 | 0.41 | 0.35 |
| 3.9 | 3.2 | 1.9 | 0 | 0.2 | 4-5-3-2-1 | 0.41 | 0.37 | 0.48 | 0.45 | 0.37 |
| 1.85 | 0 | 0.05 | 0.3 | NaN | 2-3-4-1 | 0.16 | 0.21 | 0.55 | 0.42 | NaN |

| | | | | | | | | | | |
|---|---|---|---|---|---|---|---|---|---|---|
| 1.4 | 0.55 | 0.6 | 0.5 | 0 | 5-4-2-31 | 0.14 | 0.24 | 0.59 | 0.52 | 0.013 |
| 0.35 | 0 | 1.05 | 1.25 | 1.3 | 2-1-3-4-5 | 0.50 | 0.62 | 0.47 | 0.61 | 0.33 |
| 1.2 | 0.2 | 0.55 | 0.1 | 0 | 5-4-2-3-& | 0.04 | 0.18 | 0.36 | 0.43 | 0.40 |
| 0.95 | 0.7 | 0.9 | 0 | 0.2 | 4-5-2-3_1 | 0.03 | 0.30 | 0.40 | 0.46 | 0.47 |
| NaN | 0 | 0.05 | 2.45 | 1.25 | 2-3-5-4 | 0.16 | 0.46 | 0.51 | 0.24 | 0.15 |
| 0 | 0.3 | 0.15 | 0.8 | 0.15 | 1-3=5-2-4 | 0.18 | 0.41 | 0.40 | 0.15 | 0.14 |
| 0.25 | 0.75 | 0.2 | 0 | NaN | 4-3-1-2 | 0.26 | 0.36 | 0.47 | 0.095 | NaN |
| 0 | 0.65 | 1.85 | 1.9 | NaN | 1-2-4-3 | 0.21 | 0.95 | 0.95 | 0.67 | NaN |

*4-inx-15(syb5414); inx-18(ok2454)* (7 embryos; 17 waves)

| **Delay** | | | | | | **Intensity** | | | | |
|---|---|---|---|---|---|---|---|---|---|---|
| Cell1 | Cell2 | Cell3 | Cell4 | Cell5 | | Cell1 | Cell2 | Cell3 | Cell4 | Cell5 |
| 0 | 0.5 | 0.95 | 1.05 | 1 | 1-2-3-5-4 | 0.29 | 0.55 | 0.60 | 0.44 | 0.26 |
| 2.65 | 0 | 0.75 | 1.6 . | 1.45 | 2-3-5-4-1 | 0.18 | 0.46 | 0.51 | 0.30 | 0.25 |
| 0 | 0.3 | 0.55 | 0.2 | 0.5 | 1-4-2-5-3 | 0.23 | 0.45 | 0.46 | 0.29 | 0.15 |
| 0.1 | 0 | 0.05 | 0 | 0.75 | 2=4-3-1-5 | 0.32 | 0.66 | 0.68 | 0.47 | 0.20 |
| 3 | 0 | 3 | 1.15 | 1.65 | 2-4-5-1=3 | 0.17 | 0.40 | 0.44 | 0.28 | 0.05 |
| 0.55 | 0 | 1.5 | 1.85 | 1.4 | 2-1-5-3-4 | 0.34 | 0.32 | 0.42 | 0.33 | 0.20 |
| 1.5 | 2.6 | 0 | 4 | 0.75 | 3-5-1-2-4 | 0.20 | 0.17 | 0.19 | 0.16 | 0.08 |
| 0.6 | 0.95 | 0.35 | 0 | 0 | 4=5-3-1-2 | 0.05 | 0.21 | 0.19 | 0.20 | 0.08 |
| 0.3 | 0 | 0.45 | NaN | NaN | 2-1-3 | 0.23 | 0.30 | 0.29 | NaN | NaN |
| 2.3 | 0 | 1.4 | 2.45 | 1.55 | 2-3-5-1-4 | 0.09 | 0.27 | 0.16 | 0.10 | 0.02 |
| 0.15 | 0 | 0.65 | 0.85 | 4 | 2-1-3-4-5 | 0.10 | 0.27 | 0.18 | 0.05 | NaN |
| 4.85 | 4.2 | 5.65 | 0.1 | 0 | 5-42-1-3 | 0.76 | 0.93 | 0.55 | 0.42 | 0.14 |
| 0 | 0.05 | 0.35 | 0.3 | 0.9 | 1-2-3-4-5 | 1.07 | 0.57 | 0.32 | 0.32 | 0.13 |
| 0.5 | 0.3 | 0 | 0.15 | 0.5 | 3-4-2-5=1 | 1.16 | 0.53 | 0.56 | 0.42 | 0.19 |
| 1.2 | 0 | 0.65 | 0.7 | 0.9 | 2-3-4-5-1 | 0.13 | 0.27 | 0.35 | 0.20 | 0.11 |
| 0.45 | 0.25 | 0.05 | 0 | 2.35 | 4-3-2-1-5 | 0.17 | 0.26 | 0.25 | 0.11 | 0.03 |
| 0.2 | 0.2 | 0.1 | 0 | 1 | 4-3-2=1-5 | 0.16 | 0.31 | 0.31 | 0.13 | 0.02 |

*5-deg-1(u506)* (8 embryos; 11 waves)

| Delay | | | | | | Intensity | | | | |
|---|---|---|---|---|---|---|---|---|---|---|
| Cell1 | Cell2 | Cell3 | Cell4 | Cell5 | | Cell1 | Cell2 | Cell3 | Cell4 | Cell5 |
| 4.5 | 0 | 1.65 | 1 | 4.15 | 2-4-3-5-1 | 1.03 | 1.11 | 0.44 | 0.43 | 0.58 |
| 0.65 | 0 | 0.5 | 2 | 0.9 | 2-3-1-5-4 | 0.25 | 1.73 | 1.94 | 0.91 | 0.85 |
| 2.55 | 2.7 | 0 | 1.05 | 8.7 | 3-4-2-1-5 | 0.64 | 0.69 | 2.24 | 1.73 | 0.55 |
| 6.2 | 8 | 6.2 | 2.3 | 0 | 5-4-3=1-2 | 0.70 | 1.75 | 2.94 | 0.90 | 0.49 |
| 2.85 | 2.6 | 0.5 | 5.35 | 0 | 5-3-2-1-4 | 0.60 | 2.54 | 1.31 | 0.81 | 0.52 |
| 2.15 | 2.45 | 0.2 | NaN | 0 | 5-3-1-2 | 0.46 | 2.15 | 0.98 | NaN | 0.38 |
| 0 | 0.55 | 1.85 | 0 | 0 | 1=4=5-2-3 | 0.86 | 0.58 | 1.26 | 0.74 | 0.86 |
| 3.65 | 4.3 | 3.45 | 2.55 | 0 | 5-4-3-1-2 | 0.19 | 1.08 | 0.81 | 1.39 | 0.52 |
| 0.45 | 1.35 | 0 | 1.75 | 3.15 | 3-1-2-4-5 | 0.69 | 0.91 | 1.24 | 0.48 | 0.49 |
| 1.4 | 0 | 0 | 1.05 | 1.2 | 2=3-4-5-1 | 0.07 | 1.17 | 0.28 | 0.34 | 0.72 |
| 0 | 0.15 | 2.65 | 2.65 | 0.85 | 1-2-5-3=4 | 0.40 | 0.41 | 1.60 | 1.55 | 1.22 |

*6-unc-105(n506)* (8 embryos; 12 waves)

| Delay | | | | | | Intensity | | | | |
|---|---|---|---|---|---|---|---|---|---|---|
| Cell1 | Cell2 | Cell3 | Cell4 | Cell5 | | Cell1 | Cell2 | Cell3 | Cell4 | Cell5 |
| 1 | 0.45 | 0 | 0 | 0.05 | 3=4-5-2-1 | 1.01 | 2.05 | 1.72 | 0.41 | 0.21 |
| 0 | 0.7 | 0.35 | 0.7 | 1.1 | 1-3-2=4-5 | 0.50 | 0.69 | 1.40 | 1.11 | 0.55 |
| 0 | 0.45 | 0.8 | 1 | 0.35 | 1-5-2-3-4 | 0.59 | 3.28 | 2.11 | 0.54 | 0.07 |
| 0 | 0.65 | 0.7 | 0.55 | 0.55 | 1-4=5-2-3 | 1.57 | 2.14 | 2.62 | 1.80 | 0.93 |
| 0.7 | 0 | 0.45 | 0.75 | 0.75 | 2-3-1-4=5 | 0.31 | 2.66 | 1.78 | 0.88 | 0.66 |
| 0.25 | 0 | 0.35 | 0.75 | 0.85 | 2-1-3-4-5 | 0.70 | 2.33 | 1.48 | 0.91 | 0.67 |
| 0.4 | 0.6 | 0.4 | 0 | 0.6 | 4-1=3-2=5 | 0.99 | 1.76 | 2.06 | 1.85 | 0.61 |
| 0 | 0 | 0.15 | 0.15 | 0.7 | 1=2-3=4-5 | 1.06 | 1.63 | 2.08 | 2.01 | 0.75 |
| 0 | 0.95 | 3 | 0.4 | 6.5 | 1-4-2-3-5 | 1.83 | 1.73 | 1.42 | 0.94 | 0.40 |
| 1.85 | 0 | 1.05 | 1.15 | NaN | 2-3-4-1 | 1.50 | 5.32 | 3.74 | 0.92 | NaN |
| 0.1 | 0.15 | 0.05 | 0.25 | 0 | 5-3-1-2-4 | 2.80 | 4.10 | 4.33 | 2.77 | 0.78 |
| NaN | 0.05 | 0 | 0.8 | 0.35 | 3-1-5-4 | NaN | 2.21 | 4.39 | 3.50 | 4.11 |

**Table S2:** List of the genes tested in the RNAi screen

| Gene list | Homology | Phenotype |
|---|---|---|
| *inx-12* | Innexin12 | Normal |
| *inx-13* | Innexin13 | Normal |
| *inx-14* | Innexin14 | Normal |
| *inx-15* | Innexin15 | muscle cell morphology abnormal |
| *inx-16* | Innexin16 | Normal |
| *inx-17* | Innexin17 | Normal |
| *inx-19* | Innexin19 | Normal |
| *inx-20* | Innexin20 | Normal |
| *inx-21* | Innexin21 | Normal |
| *inx-22* | Innexin22 | Normal |
| *eat-5* | Innexin | Normal |
| *inx-6* | Innexin6 | Normal |
| *inx-7* | Innexin7 | Normal |
| *inx-8* | Innexin8 | Normal |
| *inx-9* | Innexin9 | Normal |
| *inx-18* | Innexin18 | bump ventral side |
| *inx-4 (che-7)* | Innexin4 | Normal |
| *inx-10* | Innexin10 | Normal |
| *inx-11* | Innexin11 | Normal |
| *inx-1* | Innexin1 | Normal |
| *inx-2* | Innexin2 | Normal |
| *inx-3* | Innexin3 | Normal |
| *inx-5* | Innexin5 | Normal |
| *unc-9* | Innexin | Normal |
| *unc-7* | Innexin | Normal |
| *mec-4* | SCNN1B (sodium channel epithelial 1 subunit beta | Normal |
| *mec-10* | SCNN1B | Normal |
| *unc-105* | SCNN1B | Normal |
| *del-1* | SCNN1B | Normal |
| *deg-1* | SCNN1B | Normal |

| | | |
|---|---|---|
| *del-10* | SCNN1B | NOT TESTED |
| *unc-8* | SCNN1B | Normal |
| *flr-1* | ASIC | Normal |
| *trp-1* | TRPC | Normal |
| *trp-3* | TRPC | NOT TESTED |
| *cup-5* | MCOLN1 | NOT TESTED |
| *gtl-1* | TRPM | Normal |
| *gtl-2* | TRPM | Normal |
| *trpa-1* | TRPA | Normal |
| *osm-9* | TRPV | Normal |
| *ocr-1* | TRPV | Normal |
| *gon-2* | TRPM | Normal |
| *trpa-2* | TRPA | Normal |
| *acd-1* | ASIC | Normal |
| *ocr-2* | TRPV | Normal |
| *spc-1* | SPTAN1 | Muscle organisation |
| *sma-1* | SPTBN5 | Normal |
| *unc-112* | FERMT1 | Hypercontracted |
| *Itr-1* | ITPR1 | Normal |

**Table S3**: P-values of the relevant comparisons for all panels

| Figure | Genotype | p-value |
|---|---|---|
| Fig. 1F | Delay in pulsing (13 embryos, 29 waves) | Cell-1 vs Cell-2 $4.10^{-4}$ (***)<br>Cell-1 vs Cell-3 $2.10^{-2}$ (*)<br>Cell-1 vs Cell-4 0.20 (ns)<br>Cell-1 vs Cell-5 0.82 (ns)<br>Cell-2 vs Cell-3 0.21 (ns)<br>Cell-2 vs Cell- 4 $9.10^{-3}$ (**)<br>Cell-2 vs Cell-5 $8.10^{-4}$ (***)<br>Cell-3 vs Cell-4 0.21 (ns)<br>Cell-3 vs Cell-5 0.02 (*)<br>Cell-4 vs Cell-5 0.15 (ns) |
| Fig. 1G | Intensity (13 embryos, 29 waves) | Cell-1vs cell-2 $2.10^{-3}$ (**)<br>Cell-1 vs cell-3 $3.10^{-4}$ (***)<br>Cell-1 vs cell-4 0.69 (ns)<br>Cell-1 vs cell-5 0.02 (*)<br>Cell-2 vs cell-3 0.34 (ns)<br>Cell-2 vs cell-4 $6.10^{-3}$ (**)<br>Cell-2 vs cell-5 $6.10^{-7}$ (*****)<br>Cell-3 vs cell-4 $7.10^{-4}$ (***)<br>Cell-3 vs cell-5 $1.10^{-7}$ (*****)<br>Cell-4 vs cell-5 $7.10^{-3}$ (**) |
| Fig. 1H | Relative peak-intensity + relative pulse-time | Cell-1 vs Cell-2 $2.\ 10^{-9}$ (*****)<br>Cell-1 vs Cell-3 $2.10^{-9}$ (*****)<br>Cell-1 vs Cell-4 0.12 (ns)<br>Cell-1 vs Cell-5 0.18 (ns)<br>Cell-2 vs Cell-3 0.67 (ns)<br>Cell-2 vs Cell-4 $2.10^{-6}$ (*****)<br>Cell-2 vs Cell-5 $2.10^{-14}$ (*****)<br>Cell-3 vs Cell-4 $1.10^{-8}$ (*****)<br>Cell-3 vs Cell-5 $4.10^{-14}$ (*****)<br>Cell-4 vs Cell-5 $2.10^{-3}$ (**) |
| Fig. 2D | cell-2/3, (n=12 embryos) versus outside (n=12 embryos); anterior cells (n=7 embryos); tails cells (n=12 embryons) | <0,0001 (****) |
| Fig. 2D | muscles cells 2&3 (n=12 embryos) vs. tails cells" (n=12 embryons) | <0,0001 (****) |
| Fig. 2D | muscles cells 2&3(n=12 embryos) vs. anterior cells (n=7 embryos) | <0,0001 (****) |
| Fig. 2E | cell-2/3, (n=8 embryos) versus outside (n=12 embryos) | 0,03 (*) |
| Fig. 3B | wt vs *inx-18(ok2454)* | Cell-1 0.35 (ns)<br>Cell-2 0.06 (ns)<br>Cell-3 0.18 (ns) |

| | | |
|---|---|---|
| | | Cell-4 0.20 (ns)<br>Cell-5 0.40 (ns) |
| Fig. 3B | wt vs *inx-18(RNAi)* | Cell-1 0.37 (ns)<br>Cell-2 9.10-3 (**)<br>Cell-3 0.12 (ns)<br>Cell-4 0.24 (ns)<br>Cell-5 0.25 (ns) |
| Fig. 3C | wt vs *inx-18(ok2454)* | Cell-1 $9.10^{-6}$ (*****)<br>Cell-2 $3.10^{-7}$ (*****)<br>Cell-3 $2.10^{-8}$ (*****)<br>Cell-4 $3.10^{-4}$ (***)<br>Cell-5 $5.10^{-3}$ (**) |
| Fig. 3C | wt vs *inx-18(RNAi)* | Cell-1 $3.10^{-3}$ (**)<br>Cell-2 $4.10^{-3}$ (**)<br>Cell-3 $3.10^{-4}$ (***)<br>Cell-4 0.07 (ns)<br>Cell-5 0.13 (ns) |
| Fig. 3F | wt vs *inx-15(syb5414)* | Cell-1 $7.10^{-2}$ (**)<br>Cell-2 0.06 (ns)<br>Cell-3 0.19 (ns)<br>Cell-4 0.34 (ns)<br>Cell-5 0.37 (ns) |
| Fig. 3F | wt vs *inx-15(RNAi)* | Cell-1 0.06 (ns)<br>Cell-2 0.21 (ns)<br>Cell-3 0.13 (ns)<br>Cell-4 0.24 (ns)<br>Cell-5 0.41 (ns) |
| Fig. 3G | wt vs *inx-15(RNAi)* | Cell-1 0.02 (*)<br>Cell-2 $7.10^{-4}$ (****)<br>Cell-3 $2.10^{-5}$ (*****)<br>Cell-4 0.01 (*)<br>Cell-5 0.06 (ns) |
| Fig. 3G | wt vs *inx-15(syb5414)* | Cell-1 $4.10^{-5}$ (*****)<br>Cell-2 $3.10^{-4}$ (****)<br>Cell-3 $7.10^{-5}$ (*****)<br>Cell-4 $8.10^{-3}$ (***)<br>Cell-5 $1.10^{-5}$ (*****) |
| Fig. 3I | wt vs *inx-18(ok2454)* | Cell-1 0.26 (ns)<br>Cell-2 0.24 (ns)<br>Cell-3 0.21 (ns)<br>Cell-4 0.07 (ns)<br>Cell-5 0.28 (ns) |
| Fig. 3I | wt vs *inx-15(syb5414)* | Cell-1 0.14 (ns)<br>Cell-2 0.16 (ns)<br>Cell-3 0.14 (ns)<br>Cell-4 0.09 (ns)<br>Cell-5 0.09 (ns) |
| Fig. 3I | *inx-18(ok2454)* vs *inx-15(syb5414)* | Cell-1 0.08 (ns)<br>Cell-2 0.09 (ns)<br>Cell-3 0.09 (ns) |

| | | |
|---|---|---|
| | | Cell-4 0.01 (*)<br>Cell-5 0.07 (ns) |
| Fig. 3J | Relative peak-intensity + relative pulse-time<br>*inx-18(ok2454)* (6 embryos; 15 waves) | Cell-1vs Cell-2 $7.10^{-4}$ (****)<br>Cell-1 vs Cell-3 $2.10^{-5}$ (*****)<br>Cell-1 vs Cell-4 0.02 (*)<br>Cell-1 vs Cell-5 0.26 (ns)<br>Cell-2 vs Cell-3 0.16 (ns)<br>Cell-2 vs Cell-4 0.17 (ns)<br>Cell-2 vs Cell-5 0.03 (*)<br>Cell-3 vs Cell-4 $8.10^{-3}$ (**)<br>Cell-3 vs Cell-5 $1.10^{-3}$ (**)<br>Cell-4 vs Cell-5 0.31 (ns) |
| Fig. 3J | Relative peak-intensity + relative pulse-time<br>*inx-15(syb5414)* (6 embryos; 14 waves) | Cell-1 vs Cell-2 $8.10^{-5}$ (*****)<br>Cell-1 vs Cell-3 $7.10^{-4}$ (****)<br>Cell-1 vs Cell-4 0.72 (ns)<br>Cell-1 vs Cell-5 $5.10^{-5}$ (*****)<br>Cell-2 vs Cell-3 0.65 (ns)<br>Cell-2 vs Cell-4 0.01 (*)<br>Cell-2 vs Cell-5 $4.10^{-9}$ (*****)<br>Cell-3 vs Cell-4 0.03 (*)<br>Cell-3 vs Cell-5 $3.10^{-8}$ (*****)<br>Cell-4 vs Cell-5 $7.10^{-4}$ (****) |
| Fig. 3L | wild type vs *inx-15(syb5414); Pelt*-2::INX-15::mCherry | 0.02 (*) |
| Fig. 3L | *inx-15(syb5414)* vs *inx-15(syb5414); Pelt*-2::INX-15::mCherry | <0.0001 (****) |
| Fig. 4A | wt vs *deg-1(u506)* (8 embryos; 11waves) | Cell-1 0.04 (*)<br>Cell-2 0.01 (*)<br>Cell-3 0.04 (*)<br>Cell-4 $6.10^{-3}$ (**)<br>Cell-5 0.17 (ns) |
| Fig. 4A | wt vs *unc-105(n506)* (8 embryos; 12 waves) | Cell-1 0.13 (ns)<br>Cell-2 0.08 (ns)<br>Cell-3 0.15 (ns)<br>Cell-4 0.40 (ns)<br>Cell-5 0.32 (ns) |
| Fig. 4B | wt vs *deg-1(u506)* (8 embryos; 11waves) | Cell-1 0.04 (*)<br>Cell-2 0.31 (ns)<br>Cell-3 0.23 (ns)<br>Cell-4 0.47 (ns)<br>Cell-5 0.28 (ns) |
| Fig. 4B | wt vs *unc-105(n506)* (8 embryos; 12 waves) | Cell-1 0.21 (ns)<br>Cell-2 0.02 (*)<br>Cell-3 0.07 (ns)<br>Cell-4 0.10 (ns)<br>Cell-5 0.23 (ns) |
| Fig. 4E | wt vs *deg-1(u506)* (8 embryos; 11 waves) | Cell-1 0.08 (ns)<br>Cell-2 $0.3.10^{-3}$ (**)<br>Cell-3 0.07 (ns)<br>Cell-4 $8.10^{-3}$ (**)<br>Cell-5 $3.10^{-3}$ (**) |

| | | |
|---|---|---|
| Fig. 4E | wt vs *unc-105(n506)* (8 embryos; 12 waves) | Cell-1 0.08 (ns)<br>Cell-2 $3.10^{-3}$ (**)<br>Cell-0.07 (ns)<br>Cell-4 $8.10^{-3}$ (**)<br>Cell-5 $3.10^{-3}$ (**) |
| Fig. 4F | Relative peak-intensity + relative pulse-time<br>*deg-1(u506)* (8 embryos; 11waves) | Cell-1 vs Cell-2 0.03 (*)<br>Cell-1 vs Cell-3 0.01 (*)<br>Cell-1 vs Cell-4 0.30 (ns)<br>Cell-1 vs Cell-5 0.22 (ns)<br>Cell-2 vs Cell-3 0.87 (ns)<br>Cell-2 vs Cell-4 0.17 (ns)<br>Cell-2 vs Cell-5 0.16 (ns)<br>Cell-3 vs Cell-4 0.11 (ns)<br>Cell-3 vs Cell-5 0.09 (ns)<br>Cell-4 vs Cell-5 0.92 (ns) |
| Fig. 4F | Relative peak-intensity + relative pulse-time<br>*unc-105(n506)* (8 embryos; 12 waves) | Cell-1 vs Cell-2 $6.10^{-3}$ (**)<br>Cell-1vs Cell-3 0.01 (*)<br>Cell-1vs Cell-4 0.69 (ns)<br>Cell-1 vs Cell-5 0.14 (ns)<br>Cell-2 vs Cell-3 0.65 (ns)<br>Cell-2 vs Cell-4 0.01 (*)<br>Cell-2 vs Cell-5 $3.10^{-5}$ (*****)<br>Cell-3 vs Cell-4 0.03 (*)<br>Cell-3 vs Cell-5 $7.10^{-5}$ (*****)<br>Cell-4 vs Cell-5 0.05 (ns) |
| Fig.S2G | *wt vs inx-15(syb5414)* | 0.42 (ns) |
| Fig.S2G | *wt vs inx-18(ok2454)* | 0.29 (ns) |
| Fig.S2G | *wt vs unc-105(n506)* | 0.51 (ns) |
| Fig. S2F | *wt vs inx-15(syb5414)* | 0.50 (ns) |
| Fig. S2F | *wt vs inx-18(ok2454)* | 0.03 (*) |
| Fig. S2F | *wt vs unc-105(n506)* | 0.03 (*) |
| Fig. S2F | *wt vs deg-1(u506)* | 0.93 (ns) |
| Fig. S2I | Integral normalised intensity (a.u.)<br>wt vs *inx-18(ok2454)* (6 embryos; 15waves) | cell-1: 0.02 (*)<br>cell-2: $6.10^{-3}$ (**)<br>cell-3: $1.10^{-3}$ (**)<br>cell-4: $7.10^{-3}$ (**)<br>cell-5: 0.01 (*) |
| Fig. S2I | Integral normalised intensity (a.u.)<br>wt vs *inx-15(syb5414)* (6 embryos; 14 waves) | cell-1: 0.04 (*)<br>cell-2: 0.06 (ns)<br>cell-3: $6.10^{-3}$ (**)<br>cell-4: 0.03 (*)<br>cell-5: 0.02 (*) |
| Fig. S2J | wt (13 embryos, n 29 waves) vs *inx-15(syb5414); inx-18(ok2454)* (7 embryos; 17 waves) | cell1: 0.22 (ns)<br>cell2: 0.10 (*)<br>cell-3: 0.06 (ns)<br>cell-4: 0.13 (ns)<br>cell-5: 0.16 (ns) |
| Fig. S2J | *inx-15(syb5414)* vs *inx-15(syb5414); inx-18(ok2454)* | cell 1: 0.02 (*)<br>cell-1: 0.02 (*) |

| | | |
|---|---|---|
| | | cell-2: 0.44 (ns)<br>cell-3: 0.19 (ns)<br>cell-4: 0.32 (ns)<br>cell-5: 0.29 (ns) |
| Fig. S2J | *inx-18(ok2454)* vs *inx-15(syb5414); inx18(ok2454)* | cell-1: 0.42 (ns)<br>cell-2: 0.49 (ns)<br>cell-3: 0.19 (ns)<br>cell-4: 0.38 (ns)<br>cell-5: 0.10 (ns) |
| Fig. S2K | wt (13 embryos, n 29 waves) vs *inx-15(syb5414); inx-18(ok2454)* (7 embryos; 17 waves) | cell-1: $3.10^{-3}$ (**)<br>cell-2: $2.10^{-8}$ (*****)<br>cell-3: $3.10^{-10}$ (*****)<br>cell-4: $3.10^{-6}$ (*****)<br>cell-5: $3.10^{-6}$ (*****) |
| Fig. S2K | *inx-15(syb5414)* vs *inx-15(syb5414); inx-18(ok2454)* | cell-1: 0.36 (ns)<br>cell-2: $8.10^{-3}$ (**)<br>cell-3: $4.10^{-3}$ (**)<br>cell-4: 0.05 (ns)<br>cell-5: 0.34 (ns) |
| Fig. S2K | *inx-18(ok2454)* vs *inx-15(syb5414); inx-18(ok2454)* | cell-1: 0.22 (ns)<br>cell-2: 0.33 (ns)<br>cell-3: 0.02 (*)<br>cell-4: 0.04 (*)<br>cell-5: 0.01 (*) |
| Fig. S3D | Integral normalised intensity (a.u.)<br>wt vs *deg-1(n506)* (8 embryos; 11 waves) | cell-1 0.04 (*)<br>cell-2 0.03 (*)<br>cell-3 0.02 (*)<br>cell-4 0.06 (ns)<br>cell-5 0.10 (ns) |
| Fig. S3D | Integral normalised Intensity(au)<br>wt vs *unc-105(n506)* (8 embryos; 12 waves) | cell-1 0.91 (ns)<br>cell-2 0.01 (*)<br>cell-3 0.14 (ns)<br>cell-4 0.14 (ns)<br>cell-5 0.25 (ns) |

P values. *, P≤0.05; **, P≤0.01; ***, P≤0.001; ****, P≤0.0001; *****, P≤$10^{-4}$.

Tables S4 : strains used in this study

| | |
|---|---|
| *ML2143* | *(mcIs75[myo-3p::gfp::PH ; stIs10088[hlhp::his-24::mCherry; unc-119(+)]).* |
| ML2940 | *inx-15(syb5414) I* |
| RB1896 | *inx-18 (ok2454) IV* |
| HBR4 | *goeIs3[pmyo-3::GCamP3.35::unc-54–3'utr, unc-119(+)] V* |
| N2 | *Bristol* |
| ML2946 | inx-18 (ok2454) IV ; goeIs3[pmyo-3::GCamP3.35::unc-54–3'utr, unc-119(+)]V |
| ML2935 | mcIs1018(deg-1 ::GFP(syb5346) )X |
| TU1366 | deg-1 (u506) X |
| ML2944 | *deg-1 (u506) X;goeIs3[pmyo-3::GCamP3.35::unc-54–3'utr, unc-119(+)] V* |
| CZ22703 | Pmyo-3-PH-miniSOG/Pmyo-3-mCherry (juEx6916) |
| ZW295 | *inx15p::INX15;::GFP* |
| ML2936 | mcEx1017pelt-2::inx-15::mcherry::unc-54–3'utr |
| ML2943 | *unc-105(n506) II,goeIs3[pmyo-3::GCamP3.35::unc-54–3'utr, unc-119(+)] V* |
| MT1098 | unc-105(n506) II. |
| ML2935 | Deg-1::GFP<br>PHX4251 deg-1(syb4251),deg1::GFP |
| ML2934 | Unc-105::GFP PHX4276 unc-105(syb4276) |
| OH4887 | Inx18p::GFP |
| ML2938 | *mcIs75[myo-3p::gfp::PH ,mcEx1017pelt-2::inx-15::mcherry::unc-54–3'utr* |
| ML2950 | mcEx677[myo-3p::mCherry],unc-105(syb4276) |
| ML2947 | mcIs1018(deg-1 ::GFP(syb5346) , mcEx677[myo-3p::mCherry] |
| ML2936 | *mcEx1017pelt-2::inx-15::mcherry::unc-54–3'utr* |

| | |
|---|---|
| ML2946 | *inx-15(syb5414) I ,inx-18 (ok2454) IV* |
